# Safe delivery of AAV vectors to the liver of small weaned pigs by ultrasound-guided percutaneous transhepatic portal vein injection

**DOI:** 10.1101/2023.04.05.535660

**Authors:** Tanja Rothgangl, Martina Hruzova, Ralph Gnannt, Nadja Aeberhard, Lucas Kissling, Hiu Man Grisch-Chan, Sven Klassa, Nicole Rimann, Kim F. Marquart, Eleonora Ioannidi, Anja Wolf, Christian Kupatt, Xaver Sidler, Johannes Häberle, Gerald Schwank, Beat Thöny

**Affiliations:** University of Zurich, Institute for Pharmacology and Toxicology, Zurich, Switzerland; Department of Diagnostic Imaging, University Children’s Hospital Zurich, Zurich, Switzerland; Department of Farm Animals, Division of Swine Medicine, Vetsuisse Faculty, University of Zurich, 8057 Zurich, Switzerland; Division of Metabolism and Children’s Research Centre, University Children’s Hospital Zurich, Zurich, Switzerland; Zurich Center for Integrative Human Physiology, Zurich, Switzerland; Klinik und Poliklinik für Innere Medizin I, Klinikum rechts der Isar, Technical University Munich and German Center for Cardiovascular Research (DZHK), Munich Heart Alliance, Munich, Germany; Institute of Molecular Life Sciences, University of Zurich, Zurich, Switzerland; Neuroscience Center Zurich, Zurich, Switzerland

**Keywords:** pig, AAV vector, PCSK9 base editing, portal vein injection, gene therapy, large animal model

## Abstract

One challenge for liver-directed gene therapy is sufficient vector delivery to the target tissue while minimizing loss of the applied vector dose to other tissues. Infusion via peripheral veins is the least invasive approach; however, it results in systemic diffusion and substantial vector dilution. Here, we describe a safe and minimally invasive method to deliver adeno-associated virus (AAV) vectors to the liver of small weaned pigs by ultrasound-guided percutaneous trans-hepatic portal vein injection. 4-week-old piglets were infused with ∼2.5×10^14^ vector genomes comprising a dual-rAAV2/9 vector system with a split adenine base editor for *in vivo* inactivation of *PCSK9* to reduce LDL-cholesterol levels. Animals had no signs of discomfort and tolerated the procedure well. However, despite 45% editing of the target site with the applied adenine base editor system in cultivated porcine cells, we only found low amounts of AAV vector genomes and neither detectable transgene-expression nor successful editing in the treated pig livers. We hypothesize that rapid proliferation of pig hepatocytes caused AAV vector dilution, leading to a loss of the vectors from the nucleus, and hence insufficient base editor protein expression for achieving detectable editing rates. Nonetheless, ultrasound-guided percutaneous transhepatic injection to the portal vein is well-tolerated in piglets and has potential for human (neonate) application.

## Background

The discovery of the CRISPR-Cas9 technology revolutionized the genome editing field and opened new doors for personalized gene therapy. Further development of this technology led to the development of base editors, which allow precise conversion of transition point mutations and work efficiently in non-dividing cells, harboring great potential for *in vivo* correction of pathogenic mutations ^1,2^. The base editor technology is well established *in vitro* in cell lines and *in vivo* in mouse models, but its translation into clinics remains challenging. Major hurdles for base editing and gene therapy in general are safe vector systems and efficient delivery strategies, which minimize necessary doses and vector loss to unintended tissues ^3,4^. While lipid nanoparticle-based delivery of therapeutic mRNA has recently become a promising option for clinical purposes ^5^, adeno-associated virus (AAV)-based vectors are still the most widely used vectors for *in vivo* CRISPR-Cas9 delivery in preclinical studies ^6,7^. Recombinant AAV (rAAV) vectors are derived from non-pathogenic viruses and have been applied in many gene therapy clinical trials and are mostly well-tolerated. rAAVs are characterized by their small size, versatile tropism spectrum and their ability to deliver genetic information in a non-integrating manner as they reside episomally ^8^.

Due to the lack of genetically modified large animal models, several proof-of-principle genome editing therapy studies have been conducted in wildtype animals^9–12^. An attractive option for such studies is the installation of loss-of-function mutations in proprotein convertase subtilisin/kexin type 9 (PCSK9). PCSK9 is an antagonist to the low-density lipoprotein cholesterol (LDL-C) receptor and steadily counteracts LDL-C uptake. Loss-of-function mutations therefore lead to reduced LDL-C levels, which can be easily measured in the serum of animals. In addition, these mutations are associated with reduced risks for atherosclerotic cardiovascular disease, and downregulation of PCSK9 is therefore also a potential approach to treat patients with hypercholesterolemia^13–20^.

Gene therapy experiments in mouse models are indispensable for preclinical research, but studies in large animal models such as dogs, pigs or non-human primates (NHP) are necessary to ensure safety of such a therapeutic approach before its application in humans. Besides application in rodents (for a recent overview see Newby and Liu, 2021^21^), systemic injection of genome editors via AAV vectors has successfully been applied in large animals such as NHPs or pigs ^22,23^. Of note, especially the liver tissue of the porcine large animal model shows great similarities to humans in terms of weight, immune system and physiology ^24^. Neutralizing antibodies against the AAV capsid or the transgene are usually seen to be less of a concern in the murine animal model ^25^, however, such pre-existing immunity can be particularly pronounced in large animal models and can hamper the efficiency of AAV-mediated Cas9 delivery. While NHP are actually the source of many AAV serotypes, the presence of such antibodies could be less prominent in pigs ^25–27^. These advantages underscore that pre-clinical studies with pigs may support the development of novel clinically relevant treatment strategies for monogenic liver diseases.

A primary challenge when working with large animal models such as the pig (but also with human patients) is the large-scale manufacturing of vectors in high quality and purity. Preferentially, doses should be adequately chosen in order to achieve a desired effect without risking immune responses or even acute toxicity in particular with systemic injection. The standard method to deliver rAAV vectors to the liver for experimental treatment of mice, NHP and also humans, is the systemic injection, which takes advantage of the blood circulation system where most of the vector naturally ends up in the liver. Nonetheless, some viral vectors infect other organs and, although the lost amounts might be minimal, it could induce unwanted side effects, making direct vector delivery to the liver a more suitable approach. While mice tolerate rAAV vector infusion with high transduction efficacy, large animal models such as pigs and NHP exhibit severe toxicity upon systemic high-dose administration of rAAV, including systemic and local immune responses. Various methods for cell, viral vector or (naked-) DNA infusion directly to the liver were reported for pigs, including open surgery, endoscopic, and/or ultra-sound guided procedures (see: Kamimura et al., 2014; Stoller et al., 2015; Kumbhari et al., 2018; Nicolas et al., 2018; Hickey et al., 2019; Huang et al., 2021; Chan et al., 2022 ^28–34^). These direct delivery methods facilitate direct injection of vectors into the liver, which can enable higher infection rates at lower vector doses.

Here, we describe a minimally invasive injection method for the direct delivery of rAAV2/9 vectors to the liver of small weaned pigs by adopting the technology of ultrasound-guided percutaneous transhepatic portal vein injection, with the goal of achieving *in vivo* base editing in the porcine liver with clinically viable vector doses. The here presented delivery approach was proven safe, although we could not observe successful genome editing at the targeted site. This was possibly due to rapid expansion of the porcine liver, which led to vector loss prior to expression of the base editor. However, ultrasound-guided percutaneous transhepatic portal vein injection might allow the design of novel pre-clinical studies in large animals for the treatment of monogenic liver diseases.

### Animal Handling and Methods

#### Animal handling

Experiments involving mice or pigs were performed in accordance with guidelines and policies of the Veterinary Office of the State of Zurich and the Swiss law on animal protection, the Swiss Federal Act on Animal Protection (1978) and the Swiss Animal Protection Ordinance (1981). All animal studies received approval from the Cantonal Veterinary Office, Zurich, and the Cantonal Committee for Animal Experiments, Zurich, Switzerland (permissions ZH159-2020 (for mice) and ZH108-2019 (for pigs)).

Mice were housed in a pathogen-free animal facility at the Institute of Molecular Health Sciences at ETH Zurich and kept in a temperature- and humidity-controlled room on a 12-hr light/dark cycle. Mice were injected with the same dual-rAAV2/9 vectors as the piglets with a dose of 3 × 10^13^ vg/kg (1.5 × 10^13^ vg/kg per vector, see Table 1) at an age of 4 weeks via tail vein injection and sacrificed for liver isolation 46 days post injection.

**Table 1.** Characteristics of pigs in the study and infusion of rAAV vector doses.

| Pig no. | Body weight on surgery day, kg | Endpoint (post-surgery) | Injected dose of rAAV, vg/kg (body weight) | Total dose for rAAV load, vg/animal |
| --- | --- | --- | --- | --- |
| 3824 | 7.3 | day 46 | --- | --- |
| 3846 | 7.4 | day 46 | --- | --- |
| 3855 | 8.8 | day 46 | $2.9 \times 10^{13}$ | $2.5 \times 10^{14}$ |
| 3859 | 7.7 | day 46 | * $3.3 \times 10^{13}$ | $2.5 \times 10^{14}$ |
| 3886 | 8.6 | day 46 | $3.0 \times 10^{13}$ | $2.5 \times 10^{14}$ |

vg, vector genomes
\* Contrast medium injected before rAAV vector injection

Three-week-old male piglets were separated from their mothers immediately after weaning and brought to a loos barn with porcine mates 7 days before surgery. At the day of surgery their body weights were between 7.7 – 8.8 kg (n = 5; see Table 1).

#### Anesthesia and interventional procedures for piglets

Pre-anesthetic medications included i.m. infusion of a mixture of ketamine (5 mg/kg), midazolam (0.2 mg/kg) and butorphanol (0.2 mg/kg). For induction, an isoflurane mask-inhalation was used, and for maintenance, an isoflurane-oxygen mixture plus i.v. lidocain (3 mg/kg/h). Post-interventional medications included meloxicam (0.4 mg/kg i.m.). Blood withdrawals were performed (post 3 hours of fasting) on a weekly bases as a pre- and post-interventional measure for a period of 6 weeks until the end of the experiment. Blood samples were used for measuring serum antibodies against rAAV2/9 neutralizing ABs, LDL-cholesterol, and PCSK9 levels. Clinical chemical parameters were measured in the Division for Clinical Chemistry and Biochemistry at the University Children’s Hospital Zurich by automated analyzer UniCel DXC600 (Beckman Coulter, Nyon, Switzerland); more details on methods used see below.

For performing the intervention, the animals were positioned in a dorsal position and on a closed-circuit anesthesia system. The main portal vein at hepatic entry was identified using a linear transducer. To perform direct portal vein injection, an 18G needle was introduced under ultrasound control through liver tissue in an area lacking any vessels other than the portal vein.

An average total volume of 0.6 ml/kg body weight with an injection speed of approximately 0.5 ml/s was applied. When reaching the portal vein, blood was withdrawn to proof intraluminal position prior to a test injection of 1 ml NaCl 0.9%. Infusion of contrast medium was performed in one piglet to demonstrate wide distribution within the liver vasculature. When this was ensured, AAV vector genomes (vg) at doses between 2.9 to 3.3 × 10^13^ vg/kg and a total of ∼2.5 × 10^14^ per piglet were injected (Table 1). Immediately thereafter, the 18G needle was removed. Animals were investigated by ultrasound for signs of bleeding at the end of the intervention.

#### Additional materials and methods are listed in the supplementary material

## Results

### Surgical intervention in small piglets for ultrasound-guided percutaneous trans-hepatic portal vein injection

In order to test the surgical procedure of ultrasound-guided percutaneous trans-hepatic portal vein injection, we dosed three animals with rAVV2/9 and/or contrast medium as summarized in Table 1. Animals were under general anesthesia and placed in supine position for ultrasound-guided percutaneous trans-hepatic portal vein injection (Figure 1A, B). The entire procedure was done under sterile conditions described in detail in the methods section and lasted between 2 -5 minutes (time point of puncture until end of injection and removal of syringe). In brief, the main portal vein at hepatic entry was searched using a linear transducer. For injection, an 18G needle was introduced upon ultrasound in an area lacking any vessels other than the portal vein (Figure 1C-E). When placing the needle within the lumen of the portal vein, a minimal amount of blood was withdrawn prior to test injection of 1 ml NaCl 0.9%. In order to confirm successful injection and distribution, one animal was injected with contrast medium first (see Figure 1F-H) and thereafter, three animals were injected by ultrasound guidance before the needle was removed (Table 1). All piglets that underwent this procedure showed no signs of discomfort and tolerated the intervention well.

**Figure 1.**
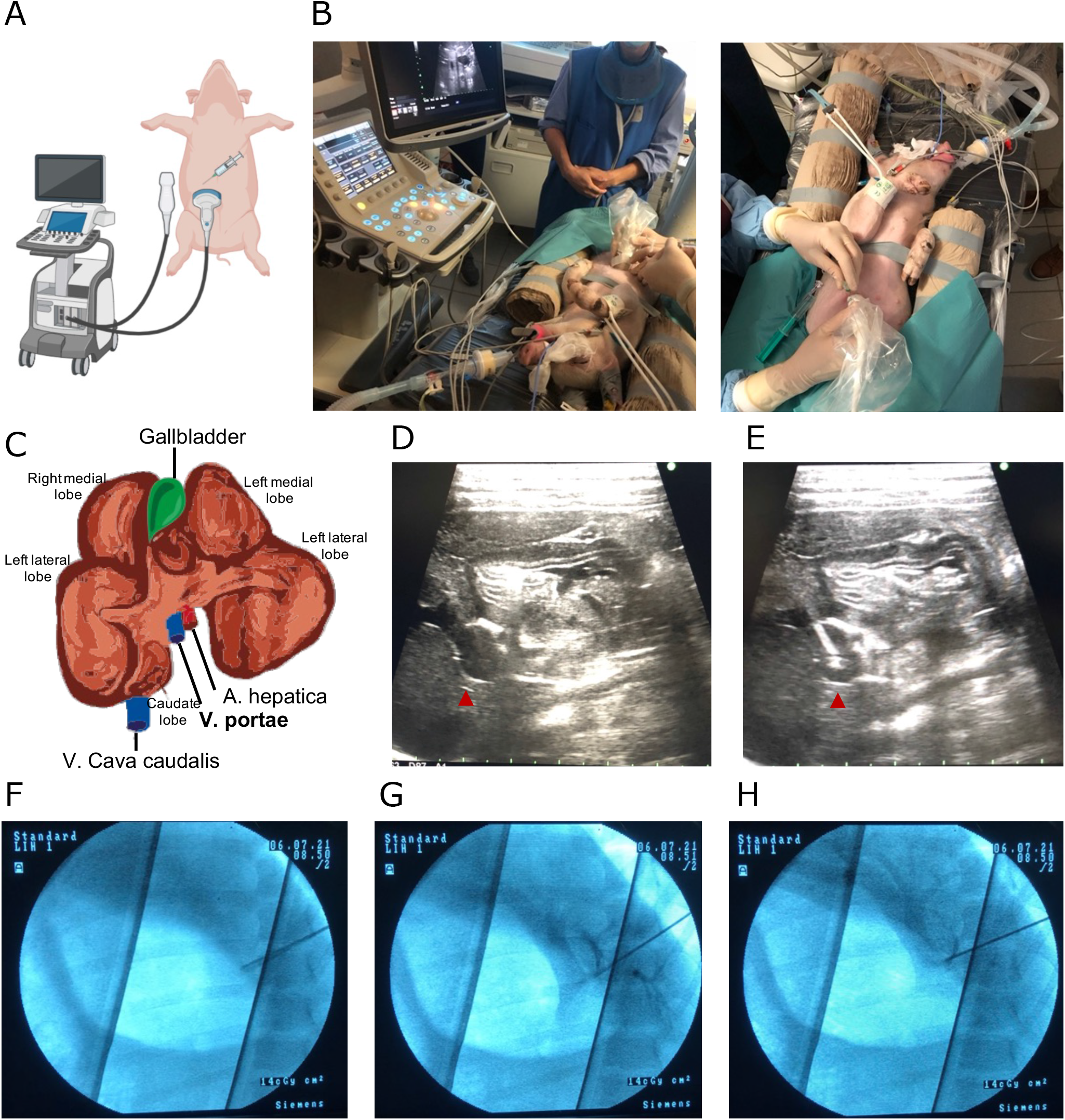
Schematics of the set-up for ultrasound-guided percutaneous trans-hepatic portal vein injection, including real-time ultrasound and X-ray examination. (A) Scheme of the experimental setup. The piglets were placed in supine position for ultrasound-guided percutaneous trans-hepatic portal vein injection using 18G needle. A saline solution (with or without rAAV2/9 vector; see Table 1) was delivered directly to the intrahepatic portal vein using a trans-hepatic access. (B) Photographs of the actual surgery setup showing the anaesthetized piglet in supine position and the ultrasound set-up for detection of the intrahepatic portal vein. (C) Illustration depicting the porcine liver and the position of the portal vein (Vena portae, in bold letters). (D-E) Real-time ultrasound-guided positioning of the needle (D) before and (E) after penetrating the portal vein (red arrow heads indicate intra-hepatic portal vein). The lower scale indicates 0.5 cm between each mark. (F-H) X-ray examination of right upper abdomen using contrast medium to ensure correct intraportal injection. X-ray pictures were taken before injection (F) and shortly after injection demonstrating distribution of the contrast medium to the left (G) and right liver lobes (H).

### Ultrasound-guided percutaneous trans-hepatic portal vein injection is a well-tolerated surgical procedure

All 5 piglets recovered in a few minutes after this intervention with no manifestations of discomfort, independent of whether they were injected with saline or an rAAV2/9-containing solution (see Table 1). Forty-six days after surgical intervention, animals were euthanized for liver resection. Macroscopic inspection of the injection sites did not show any signs of intra-parenchymal bleeding, and H&E staining did not reveal any indication for injury (Figure 2A). Likewise, blood liver markers and inflammation markers were determined after the intervention and revealed no gross changes or differences between control- and rAAV2/9 vector-infused animals (Figure 2B-G). Aminotransferases such as alanine transaminase (ALT) and aspartate transaminase (AST) as well as C-reactive protein 16 (CRP-16) are blood biomarkers, which are usually elevated when liver damage occurred. We did not observe elevated level of all three markers in comparison to the untreated control animals, indicating that the surgical procedure did not cause any liver damage (Figure 2B-D). Likewise, inflammation markers such as interleukin 6 (IL-6), tumor necrosis factor alpha (TNF-α) and interferon gamma (IFN-γ) were either unaffected or only slightly increased two days after injection in treated animals compared to untreated controls, and went back to baseline, suggesting that the surgery as well as the delivered vector doses did not cause prolonged harm to the liver of the treated animals (Figure 2E-G). From this we conclude that the procedure of ultrasound-guided percutaneous trans-hepatic portal vein injection is well-tolerated in piglets.

**Figure 2.**
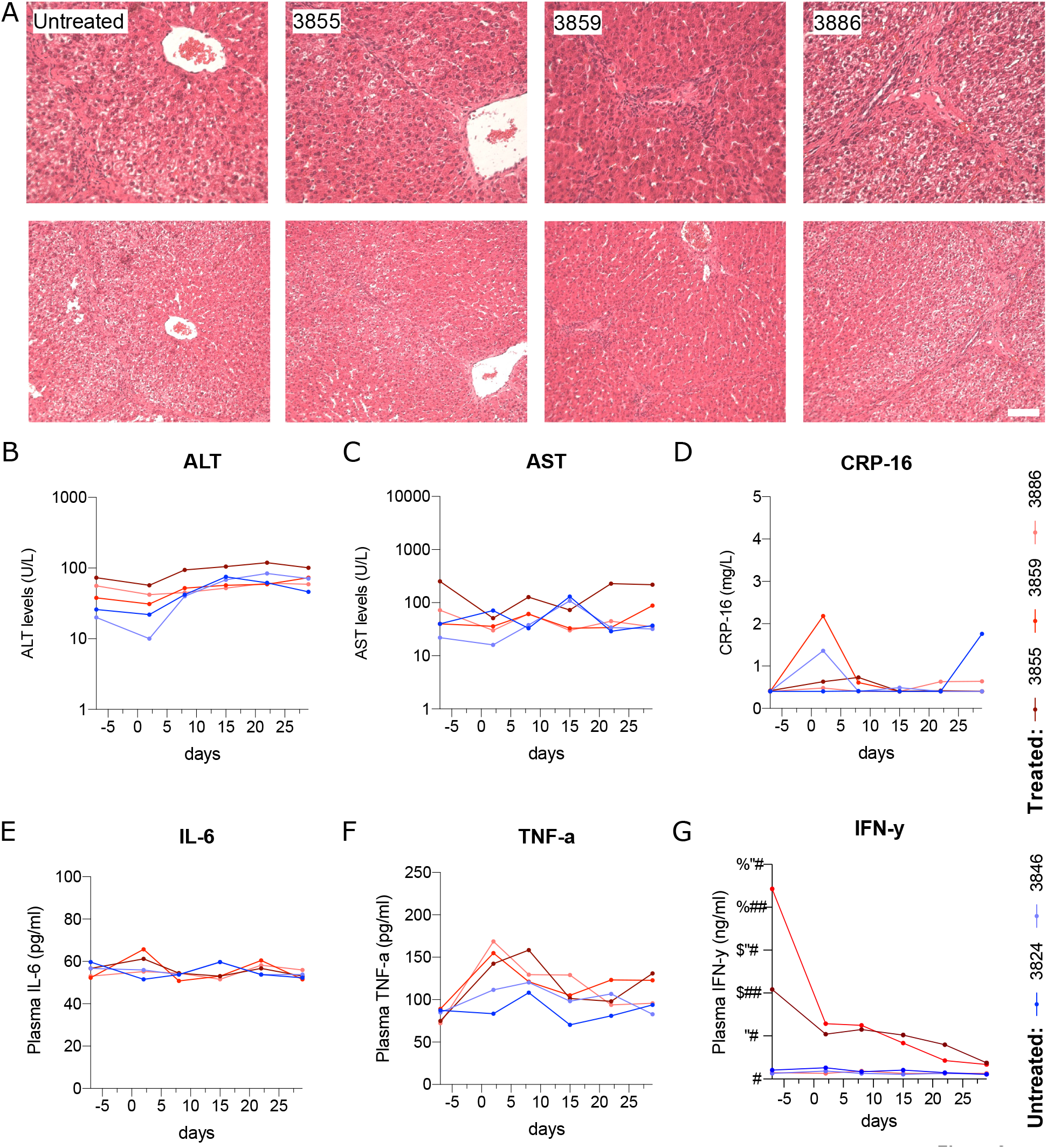
Ultrasound-guided percutaneous trans-hepatic portal vein infusion is a well-tolerated surgical procedure. (A) Representative images of hematoxylin and eosin (H&E) staining of untreated and treated pig livers. No signs of liver damage were observed. Scale bars: 50 µm (magnified view); 100 µm (overview). (B-G) Plasma levels of untreated (3824, 3846) and treated (3855, 3859, 3886) animals for (B) alanine transaminase (ALT), (C) aspartate transaminase (AST), (D) C-reactive protein 16 (CRP-16), (E) interleukin 6 (IL-6), (F) tumor necrosis factor alpha (TNF-α) and (G) interferon gamma (IFN-γ) over the experimental period of -7 days before injection at day 0 until day 29 (one month). Note that the reason for the elevated INF-γ levels in two animals prior to the intervention remained unclear and might be due to infection (diarrhea) or unnoticed injuries prior to our experiments.

### Design and validation of adenine base editing for the knockdown of porcine *PCSK9*

The three piglets (see Table 1) injected with rAAV2/9 vectors expressed an adenine base editor system that installs a loss-of-function mutation in the *PCSK9* gene ^35^. Similar to a previous study that was conducted in mice and macaques ^35^, we targeted the canonical splice-donor site of intron 1 of *PCSK9* by installing a A-to-G conversion on the reverse strand (Figure 3A). When we first transfected a plasmid expressing the adenine base editor (ABEmax^36^) and the corresponding single-guide RNA (sgRNA) into porcine LLC-PK1 kidney cells, we obtained A-to-G conversion rates at the targeted splice site of 44.8 +/-8.0% (Figure 3B, C). As the cargo capacity of AAVs is restricted to about 5 kb^37^, and full-length ABEmax together with the sgRNA does not fit on one AAV genome, we next split the base editor system in two parts. We used a previously described approach that is based on the split-intein system of *Nostoc punctiform* (Npu) to split ABEmax at the cysteine 574 of *Sp*Cas9 (the two rAAV vectors will be referred to as N-int-ABEmax and C-int-ABEmax; Figure 1D) ^35,38^. In this approach co-infection of hepatocytes will lead to the expression of N-int-ABEmax and C-int-ABEmax under the hepatocyte-specific p3 promoter^29,32^, and protein trans-splicing will lead to the reconstitution of full-length and functional ABEmax (Figure 1E)^35,38^.

**Figure 3.**
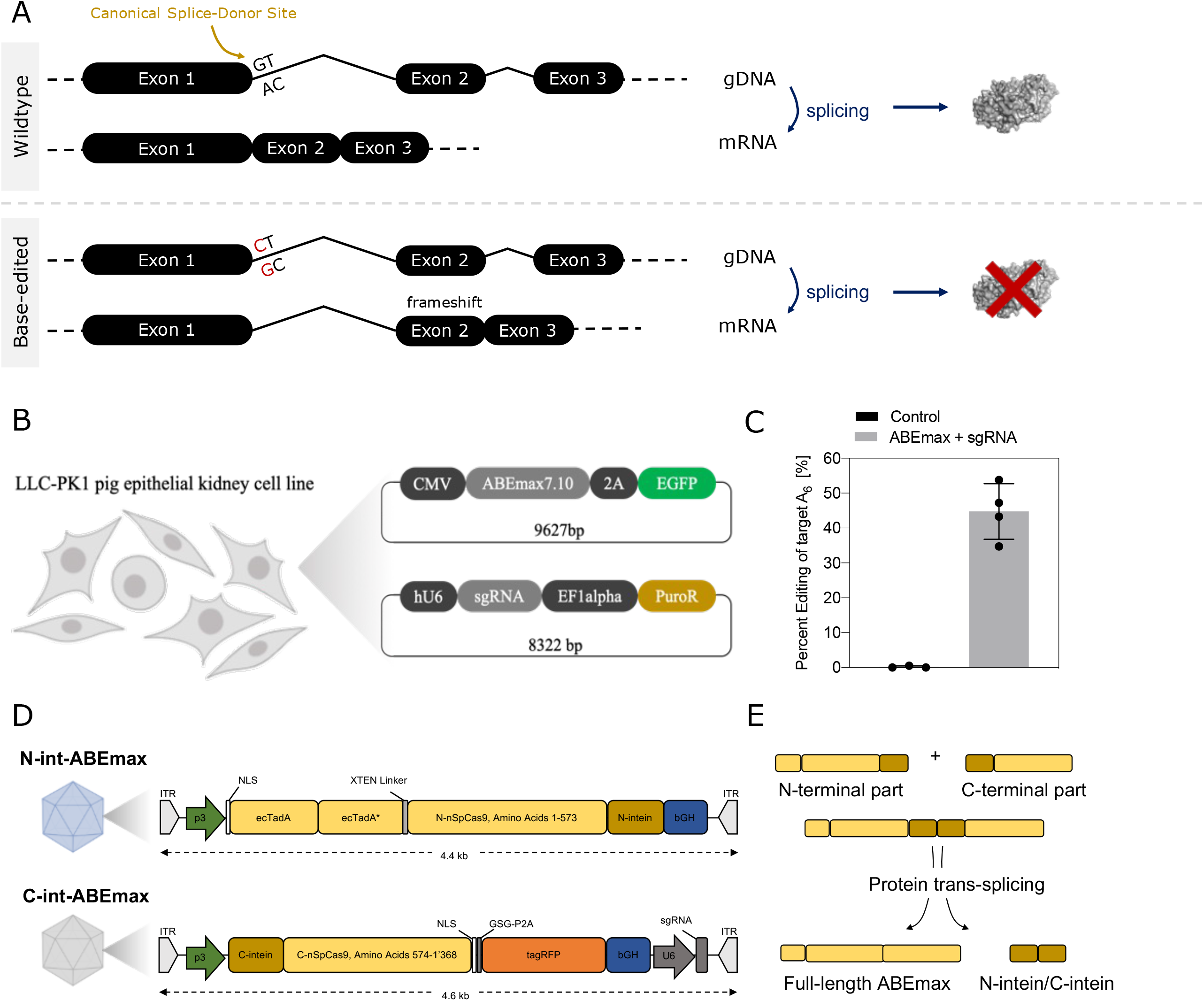
Adenine base editing approach for downregulating PCSK9. (A) Schematic of the editing approach targeting the conserved “GT” splice donor site of intron 1 in the porcine *PCSK9* gene. Sufficient editing in the genomic DNA (gDNA) leads to dysfunctional splicing of the messenger RNA (mRNA) and hence downregulation of the targeted PCSK9 protein. (B) Schematic outline of *in vitro* testing of the approach in LLC-PK1 pig kidney cells. Cells were transfected with two plasmids. One expressed ABEmax under the CMV promoter and the other expressed the targeting sgRNA under a hU6 promoter and a puromycin-resistance (PuroR) gene for selection. (C) Editing rates of the targeted adenine in the splice donor-site in transfected in LLC-PK1 pig kidney cells. Editing rates are shown for n = 3 non-targeting controls and n = 4 samples transfected with ABEmax and the targeting sgRNA. Values represent mean +/-s.d. of independent biological replicates. (D) Schematic maps of vector genomes for the dual-rAAV2/9 system. N-int-ABEmax expresses the TadA heterodimer fused to the N-terminal part of an SpCas9 nickase (nSpCas9) from the p3 promoter. C-int-ABEmax expresses the C-terminal part of nSpCas9, tagRFP, and the PCSK9 splice-site-specific sgRNA from the p3 promoter. ITR: inverted terminal repeats; NLS: nuclear localization signal; ecTadA: E. coli TadA, ecTadA*: E. coli TadA evolved; bGH: bovine growth hormone polyadenylation signal; GSG: Gly-Ser-Gly linker; P2A: porcine teschovirus-1 2A self-cleaving peptide; p3: liver-specific promoter. (E) Illustration of the applied intein-split system for the delivery of ABEmax using a dual rAAV system resulting in reconstitution of full-length ABEmax upon co-transfection of cells with both vectors and subsequent protein trans-splicing.

**Figure 4.**
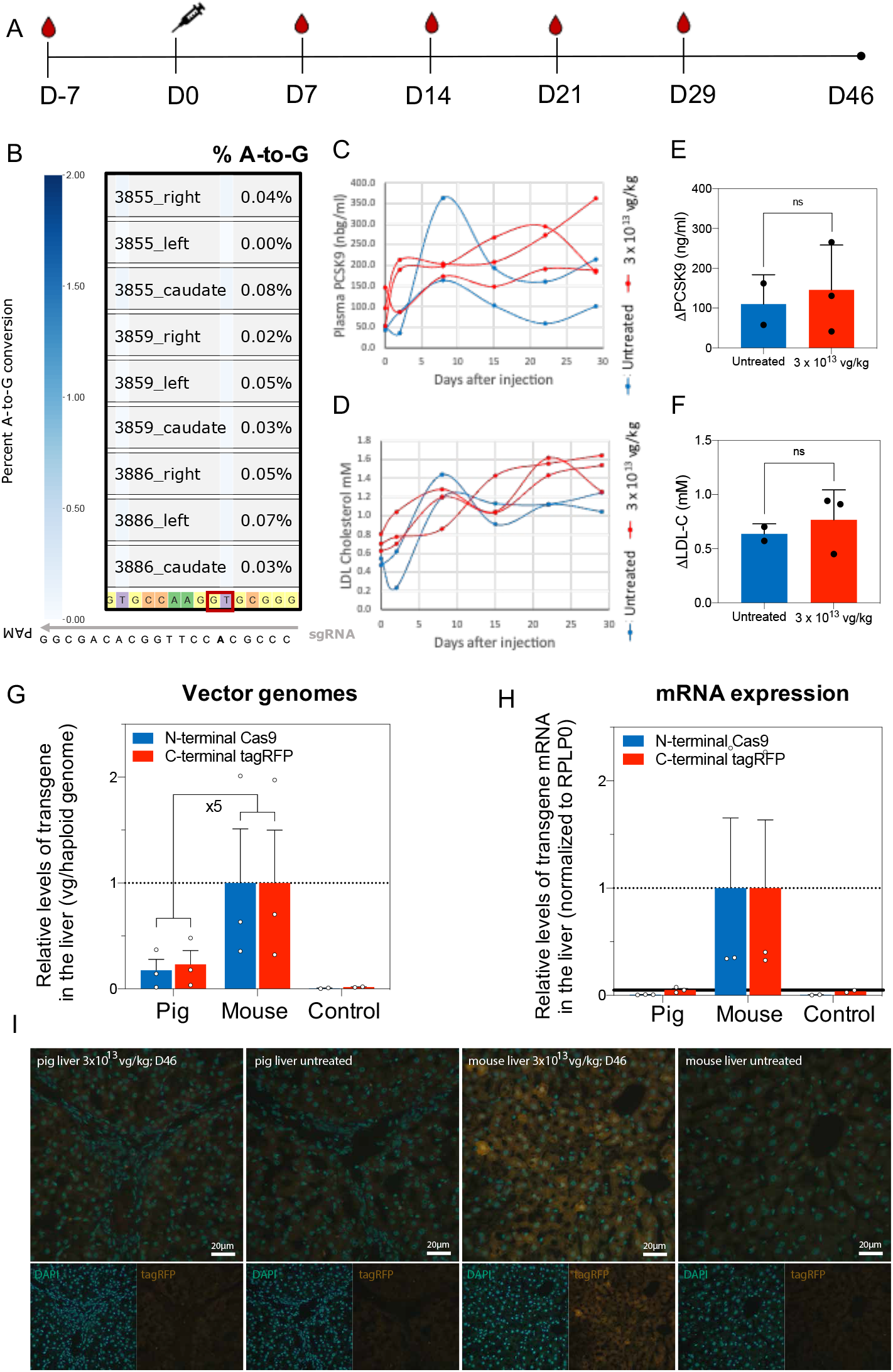
*In vivo* application for adenine base editing of the *PCSK9* gene locus in the pig liver. (A) Experimental timeline. Blood was collected (drop symbol) from all piglets right after weaning (D-7). Surgery was performed on day 0 (D0) and blood was again collected in a weekly manner for a month. The experimental time point and sacrifice of the study animals was performed on day 46 (D46). (B) Editing efficiencies on the endpoint. No A-to-G editing of the splice donor site (highlighted in red) above background was observed. (C) Plasma PCSK9 levels of untreated (blue) and treated (red) animals over time. (D) Plasma LDL-C levels of untreated (blue) and treated (red) animals over time. (E) Relative plasma PCSK9 levels compared to starting point of untreated (blue) and treated (red) animals. (F) Relative plasma LDL-C levels compared to starting point of untreated (blue) and treated (red) animals. (G) Relative levels of N-int-ABEmax and C-int-ABEmax were detected in whole liver lysates using quantitative PCR. Remaining vector genome levels of pig livers were found to be 5-times lower than in dose-matched mouse livers. (H) Relative expression levels of N-int-ABEmax and C-int-ABEmax were assayed in whole liver lysates using reverse transcription quantitative PCR. While expression of Cas9 and tagRFP was confirmed in mouse livers, vector expression levels of pig livers were below the detection limit of the assay (bold line). (I) Expression of the C-terminally-fused tagRFP, which is co-transcriptionally expressed with the C-terminal part of ABEmax under the liver-specific p3 promoter, was determined using fluorescent microscopy. Compared to dose-matched mouse livers, no tagRFP was detected in the pig livers. Scale bar: 20 µm.

### *In vivo* delivery of the adenine base editor in the pig liver using a dual-rAAV2/9 vector system

To deliver the rAAV2/9 vectors we injected them into the portal vein of three 4-week-old piglets (Table 1). Each piglet received a dose of approximately 3 × 10^13^ vg/kg and subsequently was subjected to weekly blood withdrawals for a month and was monitored until the experimental endpoint 46 days post-surgery (Figure 3A). Despite the 45% editing of the target site *in vitro* in porcine LLC-PK1 kidney cells, we did not observe any editing of the target base above background *in vivo* (Figure 3B). Monitoring PCSK9 and LDL-C levels in pig plasma over the experimental period did likewise not indicate any downregulation of PCSK9 (Figure 3C-F). Since we could exclude the presence of rAAV2/9-specific neutralizing antibodies in the serum of the piglets before injection (Suppl. Figure 1), we speculate that ABEmax was not efficiently expressed in the pigs’ hepatocytes in order to result in genome editing.

Previous studies have reported that delivery of rAAV vectors to growing or regenerating tissues can result in a loss of vector genomes and their therapeutic effect ^39^. Comparing the growth rate of 4-week-old piglets to 4-week-old mice, piglets on average double their body weight within one week while age-matched mice require 4 weeks to double their weight (Suppl. Figure 2), suggesting that the 4-week-old porcine liver proliferates proportionally faster than the murine liver. We therefore next delivered the same vector dose per kg body weight to 4-week-old juvenile mice and kept the mice for 46 days to match the experimental time-line of the pig experiment. Quantitative PCR (qPCR) for Cas9 and tagRFP on pig and mouse liver lysates revealed that pigs obtained 5-fold less vector genomes per haploid genome compared to mice (Figure 3G). Moreover, reverse transcription (RT)-qPCR to detect Cas9 and tagRFP mRNA transcript levels showed no expression above background in the liver of pigs, while expression was detectable in mice (Figure 3H). Confirming these findings, we could not observe functional tagRFP protein, which was transcriptionally co-expressed with the C-terminal part of Cas9 on the C-int-ABEmax vector, in porcine livers while expression was detected in mice (Figure 3I). Together, these data suggest that rAAV-mediated delivery of the adenine base editor into the liver of pigs likely failed because of the high proliferation rate of hepatocytes in juvenile pigs.

## Discussion

With recent advancements in genome editing therapeutic strategies for curing monogenic liver diseases have come closer to the clinics. Base editors are particularly powerful genome editing tools as they can facilitate efficient and precise correction of transition point mutations. However, before application of this technology in humans, it is of crucial importance to test safety and tolerability of vector-based genome editing strategies in large animal models.

The porcine large animal model is readily accessible as housing conditions and breeding are well established due to its extensive use in the food industry. Moreover, pigs reach sexual maturity at about eight month of age and have relatively short gestation periods with comparably large litters ^40^. Their anatomy and physiology are largely similar to humans and the model system has therefore previously been used to study surgical procedures such as organ transplantation or implantation of medical devices.

One important aspect of *in vivo* genome editing is the optimization of the delivery route to improve transduction of the target organ. In this context, we sought to investigate a delivery method for a gene therapy relevant vector by direct injection to the liver of newborn pigs with the primary focus on safety and tolerability of the intervention using genome editing as a potential readout. Here, we could show that ultrasound-guided percutaneous transhepatic portal vein injection is feasible and well-tolerated in animals. Despite the enormous amounts of rAAV achieving an average dose of 2.5 × 10^14^ vector genomes per animal, we did not observe genome editing in the liver after rAAV vector-mediated delivery of the adenine base editor. In previous studies we have shown that the here used hepatocyte-specific promoter p3 is functional in pig liver ^29,32^, and that the used ABEmax intein-split vector design allows efficient installation of splice mutations in murine *PCSK9* ^35^. We therefore hypothesize that the delivered vector dose was not sufficient to overcome the dilutive effect caused by the rapid expansion of the porcine liver. This potentially led to the loss of extrachromosomal AAV genomes from the nucleus of transduced cells and hence the absence of ABEmax protein expression. Applying faster-acting vectors, such as lipid nanoparticles (LNP) could potentially overcome this bottleneck. In contrast, the livers of 4-weeks-old (young adult) mice expand much slower (see Suppl. Figure 2), allowing second-strand synthesis of rAAV vectors and subsequent expression of the N- and C-terminal adenine base editor moieties from the p3 promoter. Future follow-up experiments with rAAV expressing e.g. a reporter gene to compare portal vein to systemic i.v. infusion might be helpful to compare these two delivery approached back-to-back and may help to estimate the necessary amount of vector or animal age required for achieving gene editing. In conclusion, we provide a minimally invasive approach to deliver rAAV, a therapeutically-relevant delivery vector, directly into the liver via the hepatic portal vein of piglets. This approach was found to be safe in the animals and compared to systemic delivery approaches could potentially reduce the required vector dose for the transduction of hepatocytes by minimizing vector loss to other organs. While, the study is lacking a back-to-back comparison to systemic injection or an injection in fully-grown animals, it nonetheless underscores certain limitations of rAAV as delivery vector and presents the applied injections route as safe and tolerable. In the future similar surgical procedures could potentially also be applied for direct liver-specific delivery of vectors for gene therapy in humans.

## Acknowledgements

We thank the VVF facility at the University of Zurich for discussions and advice as well as for providing rAAV2/9-GFP vectors for performing the neutralizing antibody assay, the University Hospital Zurich for performing immunofluorescence staining, and the Vetsuisse Faculty University of Zürich for support with pig husbandry and anesthesia. This work was supported by the Swiss National Science Foundation (CRSII5_180257 to B.T.), the PHRT grant no. 528 (to G.S., M.S., J.H. and B.T.). Development of novel therapies for liver metabolic diseases at the University Children’s Hospital Zurich is supported by the University Research Priority Program (URPP) of the University of Zurich ITINERARE (Innovative Therapies for Rare Diseases grant to JH and GS).

## Author contributions

T.R. and M.H. designed the study, performed experiments, analyzed data and wrote the manuscript together with B.T., R.G. performed percutaneous transhepatic infusion of AAV vectors to the portal vein. N.A. and X.S. were responsible for blood withdrawals, and animal handling of the piglets and crucially contributed to the logistical planning of the pig experiments. L.K., H.M.G.C, S.K., N.R., K.F.M. and E.I. performed experiments, A.W. and C.K. prepared and characterized the rAAV2/9 vectors. J.H., G.S. and B.T. designed and supervised the study and wrote the manuscript. All authors approved the final version.

## Data availability statement

All primary data are available on SRA (Sequence Read Archive) under the accession number PRJNA904094.

