## Supplementary material for "Safe delivery of AAV vectors to the liver of small weaned pigs by ultrasound-guided percutaneous transhepatic portal vein injection": Supplemetal Methods and Table

**Running title: Portal vein injection of AAV in small pigs**

Tanja Rothgangl^1*^, Martina Hruzova^1*^, Ralph Gnannt^2^, Nadja Aeberhard^3^, Lucas Kissling^1^, Hiu Man Grisch-Chan^4^, Sven Klassa^4^, Nicole Rimann^4^, Kim F. Marquart^1^, Eleonora Ioannidi^1^, Anja Wolf^6^, Christian Kupatt^6^, Xaver Sidler^3^, Johannes Häberle^4, 5, 7^, Gerald Schwank^1^, and Beat Thöny^4,5,7,8^

1 University of Zurich, Institute for Pharmacology and Toxicology, Zurich, Switzerland
2 Department of Diagnostic Imaging, University Children's Hospital Zurich, Zurich, Switzerland.
3 Department of Farm Animals, Division of Swine Medicine, Vetsuisse Faculty, University of Zurich, 8057 Zurich, Switzerland.
4 Division of Metabolism and Children’s Research Centre, University Children’s Hospital Zurich, Zurich, Switzerland
5 Zurich Center for Integrative Human Physiology, Zurich, Switzerland
6 Klinik und Poliklinik für Innere Medizin I, Klinikum rechts der Isar, Technical University Munich and German Center for Cardiovascular Research (DZHK), Munich Heart Alliance, Munich, Germany
7 Institute of Molecular Life Sciences, University of Zurich, Zurich, Switzerland
8 Neuroscience Center Zurich, Zurich, Switzerland

**Supplemental Methods**

*Cloning and generation of adeno-associated virus (AAV) vectors*

The sequences of the rAAV2/9 constructs used in this work were generated by using pLV302 and pLV312.3 (Addgene plasmid nos.119943 and 119944) where regions of interest were exchanged using NEBuilder HiFi DNA Assembly Master Mix (NEB no. E2621). Details of the inserted sequences for amino acid are listed in Suppl. Table 1. PCR was performed using Q5 High-Fidelity DNA Polymerase (New England Biolabs; Suppl. Table 2). Vector pCMV_ABEmax_P2A_GFP was a gift from David Liu (Addgene plasmid no. 112101), and LentiGuide-Puro was a gift from Feng Zhang (Addgene plasmid no. 52963). Recombinant AAV2 serotype 9 vectors were produced at the Technical University Munich and German Center for Cardiovascular Research as previously described ^1^.

*In vitro testing*

To test the *in vitro* CRISPR/Cas9 base editing we used the LLC-PK1 pig kidney cell line (ATCC CL-101). Cells were cultured in Medium 199, Earle’s Salts (ThermoFischer Scientific, 31150022) supplemented with 1% Penicillin-Streptomycin (Fischer Scientific, 15140122) and 3% fetal bovine serum at 37°C with 5% CO_2_. To test the CRISPR/Cas9 adenine base editing efficiency in LLC-PK1 cells a gRNA located in the first exon of the PCSK9 gene was chosen. Cells were transfected using the Neon Transfection System (Invitrogen) following manufacturer’s instructions. Briefly, 3 x 10^5^ cells were re-suspended with 400-450 ng of adenine base editor plasmid (pCMV_ABEmax_P2A_GFP) and 150ng of gRNA plasmid DNA (LentiGuide -Puro) in 10 μL electroporation buffer R (Invitrogen) and electroporated using the following program: 1300 V, 20 ms, 2 pulses. For the control condition we used the same ABE plasmid together with an empty gRNA plasmid. Immediately after transfection cells were transferred into 12 or 24 well cell culture dishes coated with poly-D-lysine (10 μg/mL, EMD Millipore) and cultured as usual. Transfected cells were selected 42 hrs after electroporation with puromycin (3.5 μg/mL, InvivoGen) for 3 days and harvested for gDNA extraction 7 to 10 days after transfection. Genomic DNA was extracted either with the Qiagen DNeasy Blood & Tissue Kit following manufacturer’s instructions or using the DirectPCR Lysis Reagent (Viagen). Briefly, 100 μL of DirectPCR Lysis Reagent were mixed with 5 μL of Proteinase K (Promega) and incubated for 1 hr at 55°C and then for 10 min at 85°C using the PCR thermocycler. Genomic DNA amplification and Next generation sequencing were performed as described below.

*Genomic DNA amplification, Sanger Sequencing and Next Generation Sequencing*

Tissue samples were lysed using tissue lysis buffer: (50 mM Tris-HCl at pH 8.5, 250 mM NaCl, 2.5 mM EDTA, 0.05% SDS and 1% freshly added proteinase K) and incubated at 60°C for 2 hrs and 95°C for 10 min. Target sites were amplified by polymerase chain reaction (PCR) using GoTaq G2 Hot Start Green Master Mix (Promega) or NEBNext High-Fidelity 2× PCR Master Mix (NEB BioLabs) for 30 or 26 cycles and the respective primer pair (Supplementary Table 1). Amplification products were purified using AMPure XP beads (Beckman Coulter) and sequenced with the respective in-sequence primers (Supplementary Table X) via the Sanger method to identify target locus identity. For NGS, a second amplicon was generated with primers containing sequencing adaptors for another six cycles. The products were gel purified and quantified using the Qubit 3.0 fluorometer with the dsDNA HS Assay Kit (Thermo Fisher Scientific). Samples were sequenced on Illumina MiSeq. After demultiplexing, the samples were analyzed using CRISPResso2 ^2^.

*Neutralizing Antibody Assay*

To test for the possible presence of neutralizing antibodies (NAb) against the rAAV2/9 viral vector in the pig serum we adapted a protocol from Jungmann et al. ^3^. The assay consisted of two major steps. First, the pig serum was incubated with the rAAV2/9 viral vector containing an EGFP expression marker (ssAAV2/9-CAG-EGFP-WPRE-SV40p(A), provided by the viral vector facility, UZH). Subsequently the serum-virus mixture was incubated with the HEK293T cells (ATCC CRL-321) and the signal from the expression marker was measured, which allowed to detect the presence of NAb. In detail the assay was performed as follows: HEK293T cells (5 x 10^4^ cells/well) were plated on poly-D-lysine (5 μg/mL) coated 96-well clear flat bottom plates and cultured in DMEM (Thermo Fisher Scientific) supplemented with 10% FBS and 1% P/S (Thermo Fischer Scientific) 24 hrs prior to the virus infection. Next, the pig serum was inactivated at 56°C for 35 min and incubated with the rAAV2/9 viral vector, MOI of 10,000 was used. To determine the NAb titer the following serum dilutions were tested: 1/4, 1/16, 1/64, 1/254. The viral-serum mixture was incubated for 1 hr at 37°C in DMEM supplemented with 1% P/S without FBS. Then, the viral-serum mixture was added to the HEK293T cells plated 24 hrs in advance and incubated together for 1 hr at 37°C. Two conditions were used as control, first cells cultured without serum-virus mixture and second cells cultured with virus but without the serum. After the incubation the media was replaced with DMEM supplemented with 10% FBS and 1% P/S and cells were cultured for 48 hrs. The expression of the EGFP marker was assessed using microscopy.

*Blood sampling and clinical chemistry*

Blood was collected from the external jugular vein in heparin coated tubes (Sarstedt) and centrifuged at 2,000 g for 10 min in order to gently collect plasma. Whole blood was lysed using direct lysis buffer for subsequent PCR and Sanger sequencing as described above in order to confirm that the targeted genomic site does not contain any single nucleotide polymorphisms. Liver transaminases (aspartate aminotransferase, AST; alanine aminotransferase, AST), C-reactive protein 16 (CRP-16) as well as total cholesterol, triglyceride, high-density lipoprotein (HDL) cholesterol levels were measured in the Unit for Clinical Chemistry and Biochemistry at University Children’s Hospital Zurich, Switzerland, by automated analyzer UniCel DXC600 (Beckman Coulter, Nyon, Switzerland). LDL-cholesterol was determined using the traditional Friedewald equation: total cholesterol minus HDL cholesterol minus triglycerides divided by five. PCSK9 levels were determined using Porcine PCSK9 ELISA Kit - LS-F38573-GOS15 (LSBio), interleukin 6 (IL-6) levels were determined using Porcine IL-6 Quantikine ELISA Kit - P6000B (R&D Systems), tumor necrosis factor alpha (TNF-a) levels were determined using Porcine TNF-alpha Quantikine ELISA Kit - PTA00 (R&D Systems) and Interferon gamma (IFN-y) levels were determined using Pig IFN gamma ELISA Kit - ab113353 (abcam) all according to the respective manufacturers’ instructions.

*Immunofluorescence and Histology*

Mouse and pig livers were fixed in 4% paraformaldehyde (PFA) at 4°C overnight. Tissues were transferred to a 30% sucrose solution overnight at 4°C and embedded in OCT compound in cryomolds (Tissue-Tek). Frozen tissues were sectioned at 7 µm at -20°C, and mounted directly on Superfrost Plus slides (Thermo Fisher Scientific). Cryosections were counterstained with DAPI (Thermo Fisher Scientific) and mounted in VECTASHIELD mounting medium (Vector Labs). Two frozen sections were analyzed per sample. For histological analysis, tissues were fixed using 4% PFA at 4°C overnight and dehydrated the next day before paraffinization. Paraffin blocks were cut into 5-μm-thick sections, deparaffinized with xylene and rehydrated. Sections were stained for hematoxylin and eosin (H&E) and examined for histopathological changes. Tissues were imaged using a Zeiss Axioscope for Colibri. Imaging conditions and intensity scales were matched for all images. Images were taken using Zeiss software Zen2 and analyzed by Fiji ImageJ software (v1.51n ^4^).

*Quantitative PCR*

RNA isolation was performed using the RNeasy Kit (Qiagen). cDNA was reverse transcribed using the GoScript Reverse Transcriptase Kit (Promega). RT–qPCR was performed using 5x HOT FIREPol Evagreen qPCR Supermix (Solis Biodyne) with specific primers for the delivered AAV vectors and mouse or porcine beta-actin (Suppl. Table [2](https://www.nature.com/articles/s41587-021-00933-4#MOESM1)) and analyzed by 7900HT Fast Real-Time PCR System (Applied Biosystems). Fold changes were calculated using the ΔCT method subtracting the respective value for the housekeeping gene. Vector genomes present in whole liver lysates were assessed using qPCR with genome specific or mRNA specific primers for housekeeping genes (Suppl. Table [2](https://www.nature.com/articles/s41587-021-00933-4#MOESM1)).

*Statistical analyses*

A priori power calculations to determine sample sizes for animal experiments were performed using G*Power ^5^. Statistical analyses were performed using GraphPad Prism 9.0.0 for macOS. Sample sizes and the statistical tests used are described in the figure legends. P < 0.05 was considered statistically significant.

**Supplemental Figures and Tables**

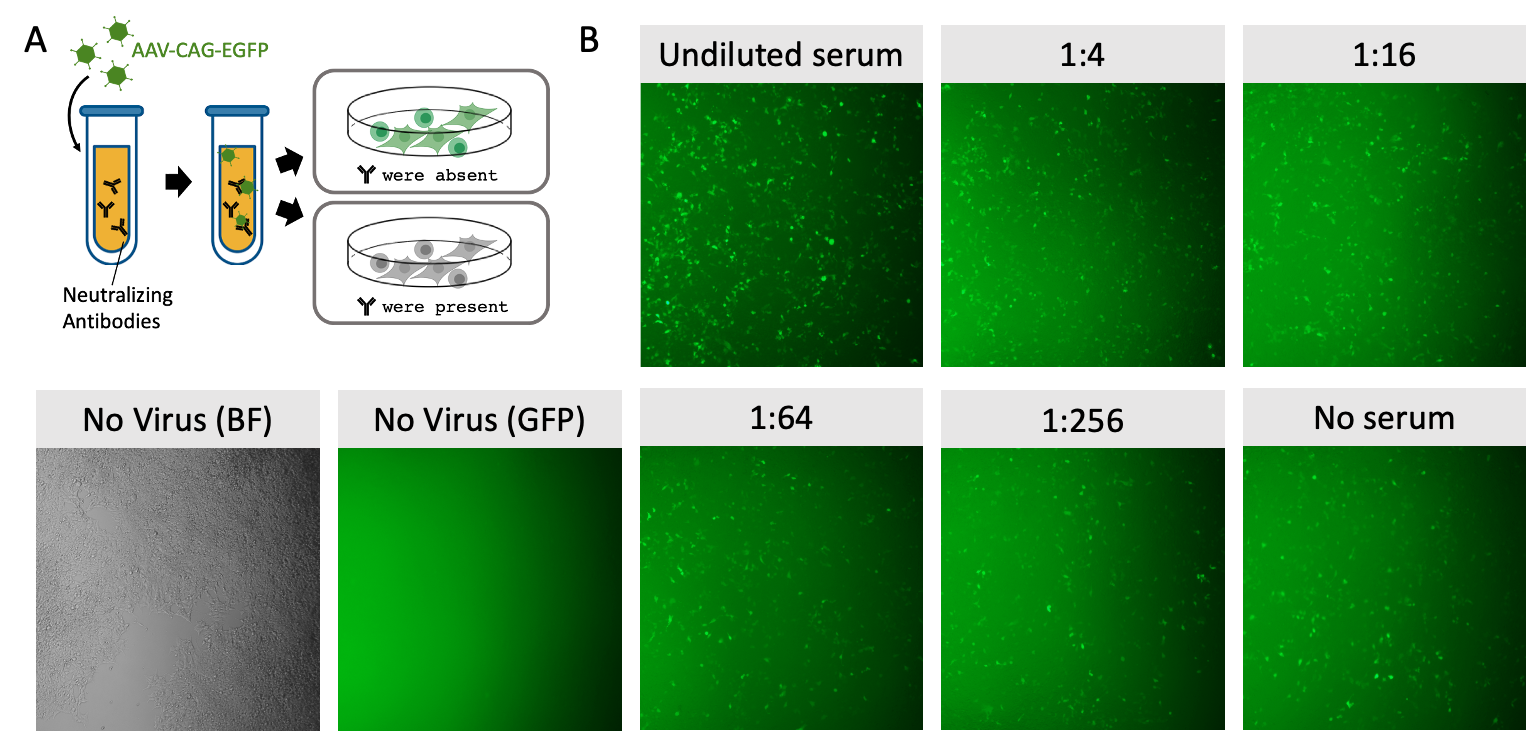

**Figure S1. Neutralizing antibody assay.** (A) Schematic illustration of the performed assay to determine neutralizing antibodies in the piglet serum against rAAV2/9. Sera were isolated from whole blood samples two weeks before the injection day and were incubated with the rAAV2/9 vectors expressing EGFP under a CAG promoter. If neutralizing antibodies are present, the AAV is unable to infect the cells, if no neutralizing antibodies are present, the cells will express EGFP. (B) Representative microscopy images showing EGFP expression after successful rAAV2/9 transduction in HEK273T cells after 48 hrs. Vector rAAV2/9 was incubated with undiluted serum and respective dilutions in PBS (1:4, 1:16, 1:64, 1:256 and PBS only). If no virus was added, no EGFP was observed. No decrease in EGFP expression upon addition of serum was observed.

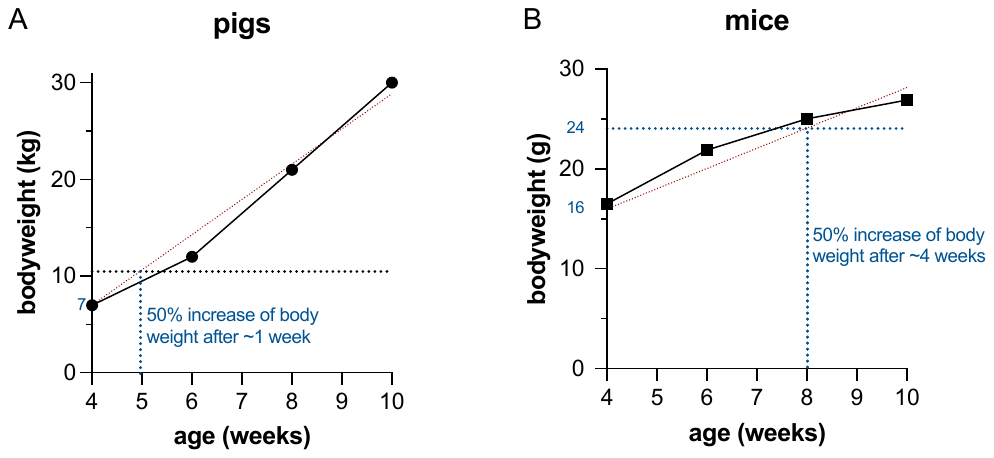

**Figure S2. Increase of bodyweight over time in pigs and mice**. (A) 4-week-old piglets reach 50% increase of body weight after ~1 week (left panel), while (B) 4-week-old mice require ~4-times longer to increase body weight by 50% (right panel). Data was obtained from (A) *Pig Growth rates - feed trough requirements*. Available at: <http://www.hendersons.co.uk/pigequip/> Pig_growth_rate.html (accessed: 16th June 2022), and from (B) *Body Weight Information for C57BL/6J*, The Jackson Laboratory. Available at: https://www.jax.org/jax-mice-and-services/strain-data-sheet-pages/body-weight-chart-000664 (accessed: 16th June 2022).

**Table S1. Details of the encoded amino acid sequences of AAV vector inserts used in this study.**

**N-int-ABEmax:** NLS - TadA-TadA* - linker - N-term-nCas9 - DnaE Npu N-intein

MKRTADGSEFESPKKKRKVSEVEFSHEYWMRHALTLAKRAWDEREVPVGAVLVHNNRVIGEGWNRPIGRHDPTAHAEIMALRQGGLVMQNYRLIDATLYVTLEPCVMCAGAMIHSRIGRVVFGARDAKTGAAGSLMDVLHHPGMNHRVEITEGILADECAALLSDFFRMRRQEIKAQKKAQSSTDSGGSSGGSSGSETPGTSESATPESSGGSSGGSSEVEFSHEYWMRHALTLAKRARDEREVPVGAVLVLNNRVIGEGWNRAIGLHDPTAHAEIMALRQGGLVMQNYRLIDATLYVTFEPCVMCAGAMIHSRIGRVVFGVRNAKTGAAGSLMDVLHYPGMNHRVEITEGILADECAALLCYFFRMPRQVFNAQKKAQSSTDSGGSSGGSSGSETPGTSESATPESSGGSSGGSDKKYSIGLDIGTNSVGWAVITDEYKVPSKKFKVLGNTDRHSIKKNLIGALLFDSGETAEATRLKRTARRRYTRRKNRICYLQEIFSNEMAKVDDSFFHRLEESFLVEEDKKHERHPIFGNIVDEVAYHEKYPTIYHLRKKLVDSTDKADLRLIYLALAHMIKFRGHFLIEGDLNPDNSDVDKLFIQLVQTYNQLFEENPINASGVDAKAILSARLSKSRRLENLIAQLPGEKKNGLFGNLIALSLGLTPNFKSNFDLAEDAKLQLSKDTYDDDLDNLLAQIGDQYADLFLAAKNLSDAILLSDILRVNTEITKAPLSASMIKRYDEHHQDLTLLKALVRQQLPEKYKEIFFDQSKNGYAGYIDGGASQEEFYKFIKPILEKMDGTEELLVKLNREDLLRKQRTFDNGSIPHQIHLGELHAILRRQEDFYPFLKDNREKIEKILTFRIPYYVGPLARGNSRFAWMTRKSEETITPWNFEEVVDKGASAQSFIERMTNFDKNLPNEKVLPKHSLLYEYFTVYNELTKVKYVTEGMRKPAFLSGEQKKAIVDLLFKTNRKVTVKQLKEDYFKKIECLSYETEILTVEYGLLPIGKIVEKRIECTVYSVDNNGNIYTQPVAQWHDRGEQEVFEYCLEDGSLIRATKDHKFMTVDGQMLPIDEIFERELDLMRVDNLPN*

**C-int-ABEmax:** DnaE Npu C-intein -C-term-nCas9, NLS-(GSG-P2A)-tagRFP

MIKIATRKYLGKQNVYDIGVERDHNFALKNGFIASCFDSVEISGVEDRFNASLGTYHDLLKIIKDKDFLDNEENEDILEDIVLTLTLFEDREMIEERLKTYAHLFDDKVMKQLKRRRYTGWGRLSRKLINGIRDKQSGKTILDFLKSDGFANRNFMQLIHDDSLTFKEDIQKAQVSGQGDSLHEHIANLAGSPAIKKGILQTVKVVDELVKVMGRHKPENIVIEMARENQTTQKGQKNSRERMKRIEEGIKELGSQILKEHPVENTQLQNEKLYLYYLQNGRDMYVDQELDINRLSDYDVDHIVPQSFLKDDSIDNKVLTRSDKNRGKSDNVPSEEVVKKMKNYWRQLLNAKLITQRKFDNLTKAERGGLSELDKAGFIKRQLVETRQITKHVAQILDSRMNTKYDENDKLIREVKVITLKSKLVSDFRKDFQFYKVREINNYHHAHDAYLNAVVGTALIKKYPKLESEFVYGDYKVYDVRKMIAKSEQEIGKATAKYFFYSNIMNFFKTEITLANGEIRKRPLIETNGETGEIVWDKGRDFATVRKVLSMPQVNIVKKTEVQTGGFSKESILPKRNSDKLIARKKDWDPKKYGGFDSPTVAYSVLVVAKVEKGKSKKLKSVKELLGITIMERSSFEKNPIDFLEAKGYKEVKKDLIIKLPKYSLFELENGRKRMLASAGELQKGNELALPSKYVNFLYLASHYEKLKGSPEDNEQKQLFVEQHKHYLDEIIEQISEFSKRVILADANLDKVLSAYNKHRDKPIREQAENIIHLFTLTNLGAPAAFKYFDTTIDRKRYTSTKEVLDATLIHQSITGLYETRIDLSQLGGDSGGSKRTADGSEFEPKKKRKVGSGATNFSLLKQAGDVEENPGPMVSKGEELIKENMHMKLYMEGTVNNHHFKCTSEGEGKPYEGTQTMRIKVVEGGPLPFAFDILATSFMYGSRTFINHTQGIPDFFKQSFPEGFTWERVTTYEDGGVLTATQDTSLQDGCLIYNVKIRGVNFPSNGPVMQKKTLGWEANTEMLYPADGGLEGRSDMALKLVGGGHLICNFKTTYRSKKPAKNLKMPGVYYVDHRLERIKEADKETYVEQHEVAVARYCDLPSKLGHKLN*

**Table S2. Oligonucleotides for PCR amplification used is this study.**

| Primer | 5’-3’ sequence |
| --- | --- |
| PCSK9_pig_NGS_F | CTTTCCCTACACGACGCTCTTCCGATCTNNNNNNAGGTCGGAGGAGGACATTCT |
| PCSK9_pig_NGS_R | GGAGTTCAGACGTGTGCTCTTCCGATCTNNNNNNGAGTAAACCGAGGACGGAGA |
| Sybr_N-term_Cas9_F | AGGATAAGAAGCACGAGCGG |
| Sybr_N-term_Cas9_R | ACTTGATCATGTGGGCCAGG |
| Sybr_tagRFP_F | AGAAAACACTCGGCTGGGAG |
| Sybr_tagRFP_R | GTCCACATAGTAGACGCCGG |
| hmlg_mus+pig_RPLP0_F | TGCTTCATTGTGGGAGCAGA |
| hmlg_mus+pig_RPLP0_R | GCATCATGGTGTTCTTGCCC |
| hmlg_RPLP0_EXspan_F | GACCTCCTTCTTCCAGGCTTT |
| hmlg_RPLP0_EXspan_R | ATCAGCTGCACATCACTCAG |
